# Spatial Transcriptomics-correlated Electron Microscopy

**DOI:** 10.1101/2022.05.18.492475

**Authors:** Peter Androvic, Martina Schifferer, Katrin Perez Anderson, Ludovico Cantuti-Castelvetri, Hao Ji, Lu Liu, Simon Besson-Girard, Johanna Knoferle, Mikael Simons, Ozgun Gokce

## Abstract

Current spatial transcriptomics methods identify cell states in a spatial context but lack morphological information. Scanning electron microscopy, in contrast, provides structural details at nanometer resolution but lacks molecular decoding of the diverse cellular states. To address this, we correlated MERFISH spatial transcriptomics with large area volume electron microscopy using adjacent tissue sections. We applied our technology to characterize the damage-associated microglial identities in mouse brain, allowing us, for the first time, to link the morphology of foamy microglia and interferon-response microglia with their transcriptional signatures.

## Main

Spatial transcriptomics (ST) methods provide spatially resolved gene expression profiling for in-depth characterization of cell types and states within a tissue^1^. These maps offer unprecedented view of cellular location and molecular phenotype but they lack the ability to resolve tissue ultrastructure. On the contrary, scanning electron microscopy (EM) provides a nanometer-resolution view of tissue ultrastructure but is limited in its capacity to directly assign molecular identities^2, 3^. To bridge molecular and morphological phenotypes, we developed **S**patial **T**ranscriptomics-correlated **E**lectron **M**icroscopy (STcEM) which correlates large-area scanning EM and multiplexed error-robust fluorescence in situ hybridization (MERFISH)^4^ and links transcriptional identities of single cells with ultrastructural data. The MERFISH is a single-molecule spatial transcriptomics technology capable of measuring hundreds to thousands of genes simultaneously with single-cell resolution^5^. This makes MERFISH a compelling ST method to integrate with EM, with the shortcoming that sample preparation requirements substantially differ from EM. The MERFISH protocol is based on snap-frozen tissue with subsequent washing, embedding and tissue clearing steps that destroy tissue ultrastructure. EM, in comparison, requires chemical fixation, heavy metal contrasting and resin embedding steps for image formation, thereby prohibiting subsequent investigation by currently available ST methods. Therefore, we harmonized both protocols to a common ground until cryo-microtomy allowed adjacent, 10 μm thin sections to be processed for EM and MERFISH, respectively (Figure 1A).

**Figure 1.**
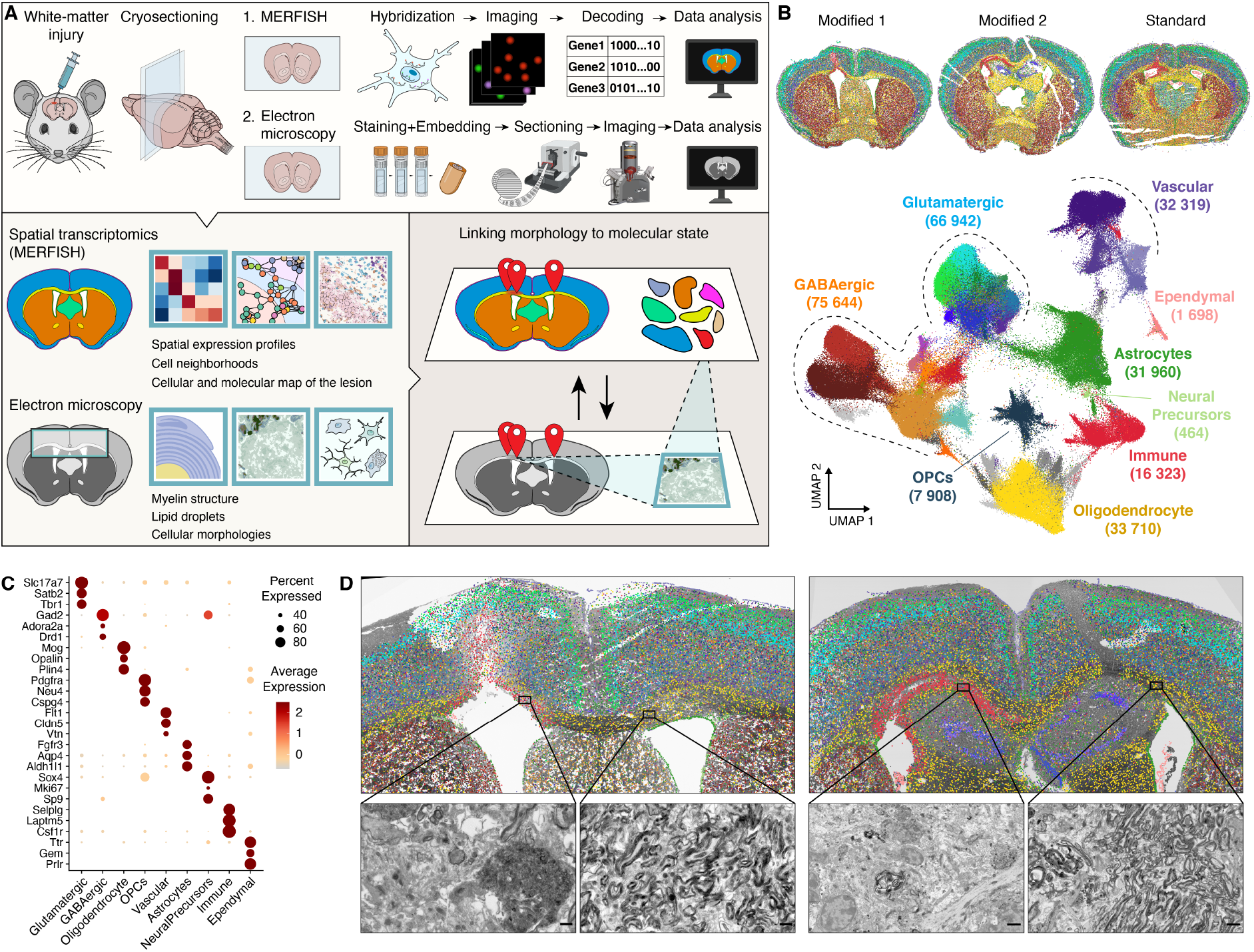
STcEM spatially links single cell transcriptomes with tissue ultrastructure. A) Overview of the STcEM method. Adjacent sections of the same sample are processed in parallel by optimized MERFISH and EM protocols and spatially aligned to directly link transcriptional profiles with nanomorphology of the regions of interest. B) Transcriptional identities of single cells with their spatial location in tissue (top) and embedded by UMAP (bottom). C) Bubble plot showing expression of cell type markers in identified single cell populations. D) MERFISH data registered onto the 2D overview EM micrograph. Zoomed-in areas show myelin structure in LPC-injected (left) and uninjured (right) white matter (corpus callosum).

We then evaluated the quality of MERFISH data relative to the standard protocol (Figure S1). Brain sections obtained with our STcEM protocol had similar segmented cell count, cell dropout rate, volume, background signal and showed excellent correlation with the standard protocol with slightly lower sensitivity (Figure S1A-C). The MERFISH analysis identified all major brain cell types including glutamatergic and GABAergic neurons, oligodendrocytes, immune cells, astrocytes, oligodendrocyte-precursor cells, ependymal and vascular cells and localized them to their original spatial positions (Figure 1B-C). Moreover, by combination of subclustering analysis and reference mapping, we identified smaller cellular subpopulations such as various classes of vascular and immune cells or layer-specific neurons in cortex, whose spatial organization matched expected patterning (Figure S2A). Importantly, these results were consistent between sections prepared with modified and standard MERFISH protocol (Figure S2B-C), altogether demonstrating the feasibility of the STcEM protocol for MERFISH. In order to generate as much complementary ultrastructural information using large-area EM, we flat-embedded the entire coronal section and generated semithin sections with a block face covering both hemispheres allowing us to scan 3×5mm^2^ area, which is at the limit of available sectioning tools (Figure 1D).

Next, we used STcEM to investigate a toxin-induced model of CNS injury, in which a single injection of lysophosphatidylcholine (LPC) (lysolecithin) induces a focal demyelinating lesion in the white matter of the mouse brain. White matter is composed of mostly myelinated axons that connect neurons from different brain regions into functional circuits. Due to the size and complexity of myelinated axons, EM is the gold standard for visualizing their structure and organization^6^. After demyelinating injury, microglia and macrophages proliferate and migrate into demyelinating lesions where they phagocytose and metabolize lipid-rich myelin, clear damaged myelin sheaths, initiate repair mechanisms and communicate with cells of the adaptive immune system^7–9^. Previous studies revealed the heterogeneity of microglial transcriptional states in different regions and conditions^10^, including disease-associated microglia (DAM)^11^, activated response microglia (ARM)^12^, lipid-droplet-accumulating microglia (LDAM)^13^, injury-responsive microglia^14^ or interferon response microglia (IRM)^12^. However, spatial as well as annotated ultrastructural information of the different microglial states in and around the lesion is lacking. Thus, we used STcEM to integrate ultrastructural changes in myelin sheaths and microglial morphology with cellular transcriptional states in demyelinating lesions. We registered MERFISH data onto EM sections to identify regions of interest where microglia and T cells respond to injury. The lesion site was apparent in MERFISH data by profound accumulation of microglia and absence of oligodendrocytes, while the contralateral hemisphere showed none of such characteristics (Figure 1D). EM micrographs showed an area with demyelinated axons and degenerated cellular debris that overlapped with the lesion area identified by MERFISH (Figure 1D).

Subclustering analysis of MERFISH data revealed the presence of four microglial clusters (Figure 2A). One homeostatic cluster, expressing high levels of typical microglial markers such as Tmem119, Csf1r, was enriched in the uninjured hemisphere. The remaining clusters localized predominantly to the injured area indicating damage associated microglial states (Figure 2B). One of these clusters was marked by upregulation of interferon-stimulated genes (Stat1, Ifit1, Rsad2) similar to previously described interferon-response microglia (IRM)^12, 14^. Another cluster was defined by upregulation of DAM markers such as Apoe, Itgax, Clec7a and was enriched in lesion site, hence we refer to it as DAM. The fourth cluster shared DAM markers, but displayed, in addition, a highly specific upregulation of Gpnmb, Lgals3 and genes related to lipid droplet formation and cholesterol metabolism such as Plin2, Soat1, Abca1 and Abcg1; and we therefore refer to this population as lipid-associated microglia. Lipid-associated microglia localized predominantly close to the ventricle edge and were enriched in the lesion core as opposed to the activated population, which was spread more equally around the entire demyelinated area (Figure 2D, S3A).

**Figure 2.**
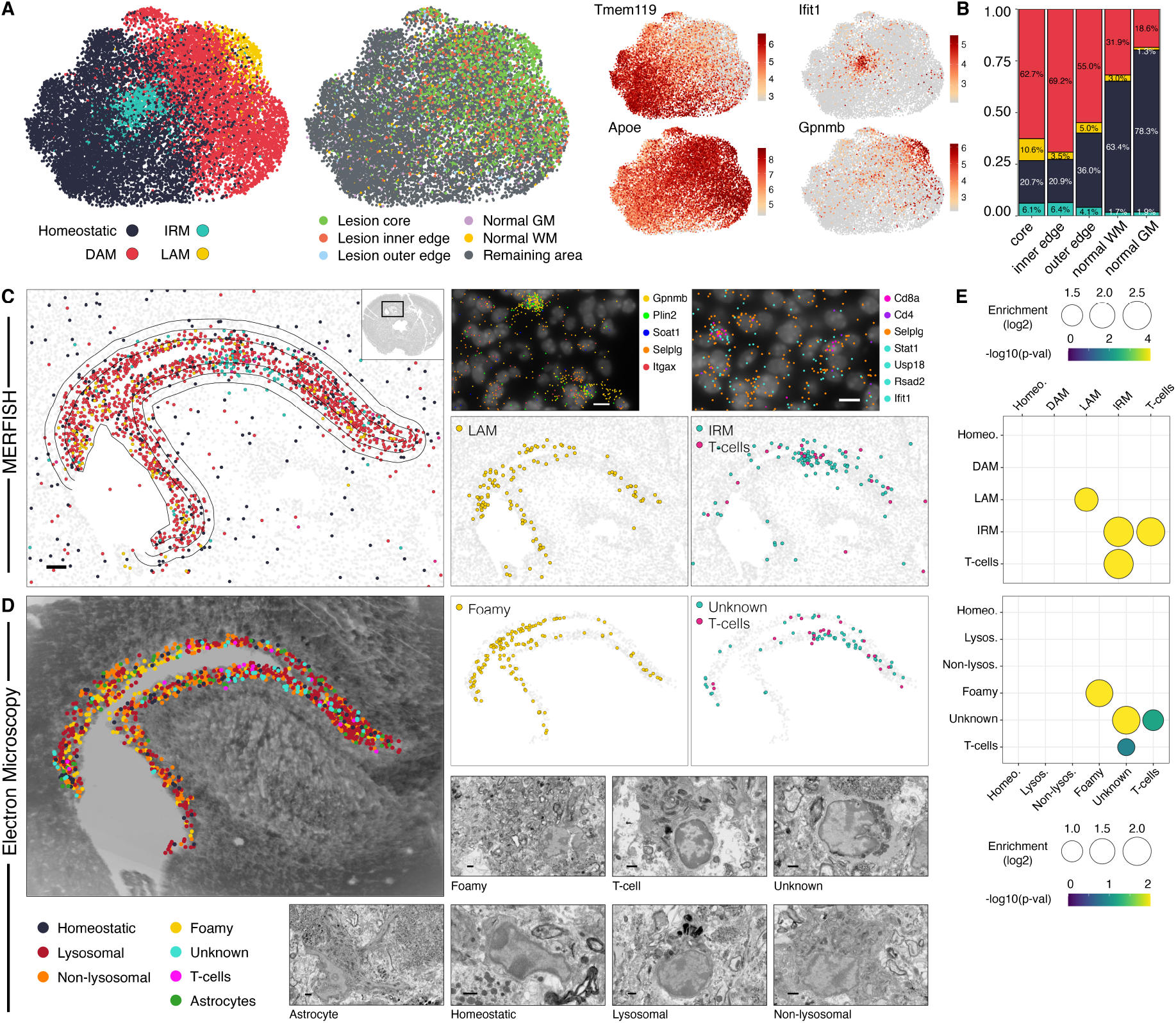
STcEM analysis of microglia and T-cells. A) UMAP plots of microglia colored by identified clusters (left), tissue region (middle), and expression of marker genes (right). DAM = disease-associated microglia, IRM = interferon-response microglia, LAM = lipid-associated microglia. B) Frequency of microglial clusters per tissue region. C) Transcriptional annotation of microglia and T-cells based on MERFISH and their spatial locations in WM lesion. Polygons depict segmented lesion areas. Zoomed-in plots (top right) show spatial location of individual transcripts of selected marker genes superimposed over DAPI signal. D) Morphological annotation of microglia and T-cells based on EM and their spatial locations in WM lesion superimposed onto a summed EM image stack. Representative EM images per each category are shown. Scale bar 1 μm. E) Neighborhood analysis of MERFISH data (top) and EM data (bottom) showing enrichment of target cell types (x-axis) among 5 nearest neighbors of query cell type (y-axis). P-values and fold-enrichment are derived from empirical distribution obtained from 10^4^ random permutations.

To further characterize and validate MERFISH-identified microglial populations on a full-transcriptome scale, we performed single-cell RNA sequencing (scRNA-Seq) on sorted CD11b+ cells using SmartSeq2 protocol in a new cohort of mice. After QC and exclusion of smaller contaminating cell populations, we obtained 1017 single microglial transcriptomes from 3 control and 5 LPC-injected mice (Figure S4A-C). Mapping MERFISH-derived labels onto scRNA-Seq data revealed the presence of equivalent microglial populations with highly concordant transcriptional signatures (Figure S3B-C). Leveraging the full-transcriptome data, we performed functional enrichment analysis focusing on lipid-associated microglia, which revealed a strong upregulation of pathways related to lipid metabolism and lysosomal processing (Figure S3D), providing further evidence that these cells are engaged in lipid-processing functions. Analyzing activity scores of published single cell signatures of lipid-associated macrophages from various tissues^15–17^ revealed that our lipid-associated microglia cluster upregulated consensus lipid-associated macrophage signature, while the remaining clusters upregulated either homeostatic, interferon or DAM signatures (Figure S3E), providing additional evidence that our lipid-associated microglia cluster represents white-matter counterpart of peripheral lipid-loaded macrophages.

To link the transcriptional and morphological phenotype of injury-responding cells, we collected serial semithin sections of the entire thickness of the adjacent vibratome section. These large area (3×5 mm^2^) sections were used to acquire volume EM data of the lesion areas at 20 nm lateral resolution. While previous light microscopy correlative studies divided tissues into chunks of 1-3 mm edge length^18^, we preserved the coronal section throughout processing which allowed uninterrupted alignment to the MERFISH images. To our knowledge this is one of the largest area scanning EM datasets complementing current large volume contrasting^19^ and fast imaging efforts^20^. Blind to the MERFISH data, we annotated cells in the demyelinated area into categories according to their ultrastructural morphology in multiple EM sections (Figure 2D). Myeloid cells were the most abundant immune cell type in the lesion center, in agreement with the microglial localization in the MERFISH data. In addition, we identified a rare population of T-cells in both MERFISH and EM data (Figure 2C-D). EM annotation revealed multiple myeloid subgroups, including normal-appearing microglia, activated myeloid cells with high content of lysosomes, myeloid cells with activated morphology but low lysosomal content, foamy myeloid cells characterized by excessive deposition of lipid droplets in their cytoplasm and one other class of unknown immune cell type (Figure 2D). Strikingly, foamy myeloid cells were spatially highly correlated with lipid-associated microglia identified by MERFISH, providing evidence that they represent the same cell state. Neighborhood analysis revealed that lipid-associated microglia are significantly more often found in the proximity of cells of the same state, often forming clusters of lipid-loaded microglia (Figure 2E top). Notably, this was in excellent agreement with the EM data, where foamy myeloid cells also clustered together (Figure 2E bottom). Spatial matching of remaining classes revealed strong concordance of patterns of homeostatic microglia, and activated microglia classes, demonstrating how intersection of morphological and molecular phenotype provided by STcEM can reveal tissue organization.

To test the ability of STcEM to assign identity to a cell of unrecognized morphology, we focused on the unknown cell type which displayed a characteristic heterochromatin pattern and perinuclearly-clustered organellar content in the cytoplasm. These cells often co-localized with T cells in EM annotations (Figure 2D-E). Their spatial distribution matched positions of IRM in adjacent MERFISH section (Figure 2C-D), where IRM also significantly co-localized with T-cells (Figure 2D-E), suggesting the cells with unrecognized EM ultrastructure are IRMs. Together, our results demonstrate how the integration of morphological and molecular phenotypes by STcEM can assign identity to unknown cell states.

In summary, STcEM extends the capacity of current EM methods such as correlative light and electron microscopy or immunogold labeling from handful of antibodies to hundreds of molecular markers simultaneously, while preserving unprecedented EM resolution. This dramatic increase in gene throughput and molecular resolution allows for unbiased identification of subtle cellular states and molecular pathways opening new opportunities to link transcriptional profiles to ultrastructure in biology and pathology. While in this study we focused on brain, STcEM can be in principle applied to any tissue.

One of STcEM’s current limitations is that using 10 μm thin adjacent sections prohibits acquisition of ST and EM data from the same cell. However, our results show that spatial niches often extend to hundreds of microns and thus we were able to connect transcriptional states to cell morphology in adjacent sections. We envision future developments, where ultrathin sectioning of the tissue could generate multiple sections from a single cell and allow identification of the individual transcripts in cellular organelles segmented from EM images, eventually providing a powerful approach for subcellular transcriptomics.

The current revolution in the generation of large volume EM datasets calls for new strategies for the annotation of cells. Currently, the cell identities in EM datasets are mostly manually annotated in an unintentionally biased way^21, 22^. Large variations between manual annotations limits the application of deep-learning approaches for automation. In its current form, STcEM improves the manual annotations by molecularly defining the cellular states in regions. In the future, STcEM may be used to decrease the required level of human supervision and generate high quality training data-sets for deep-learning approaches.

## SUPPLEMENTARY FIGURES

**Figure S1.**
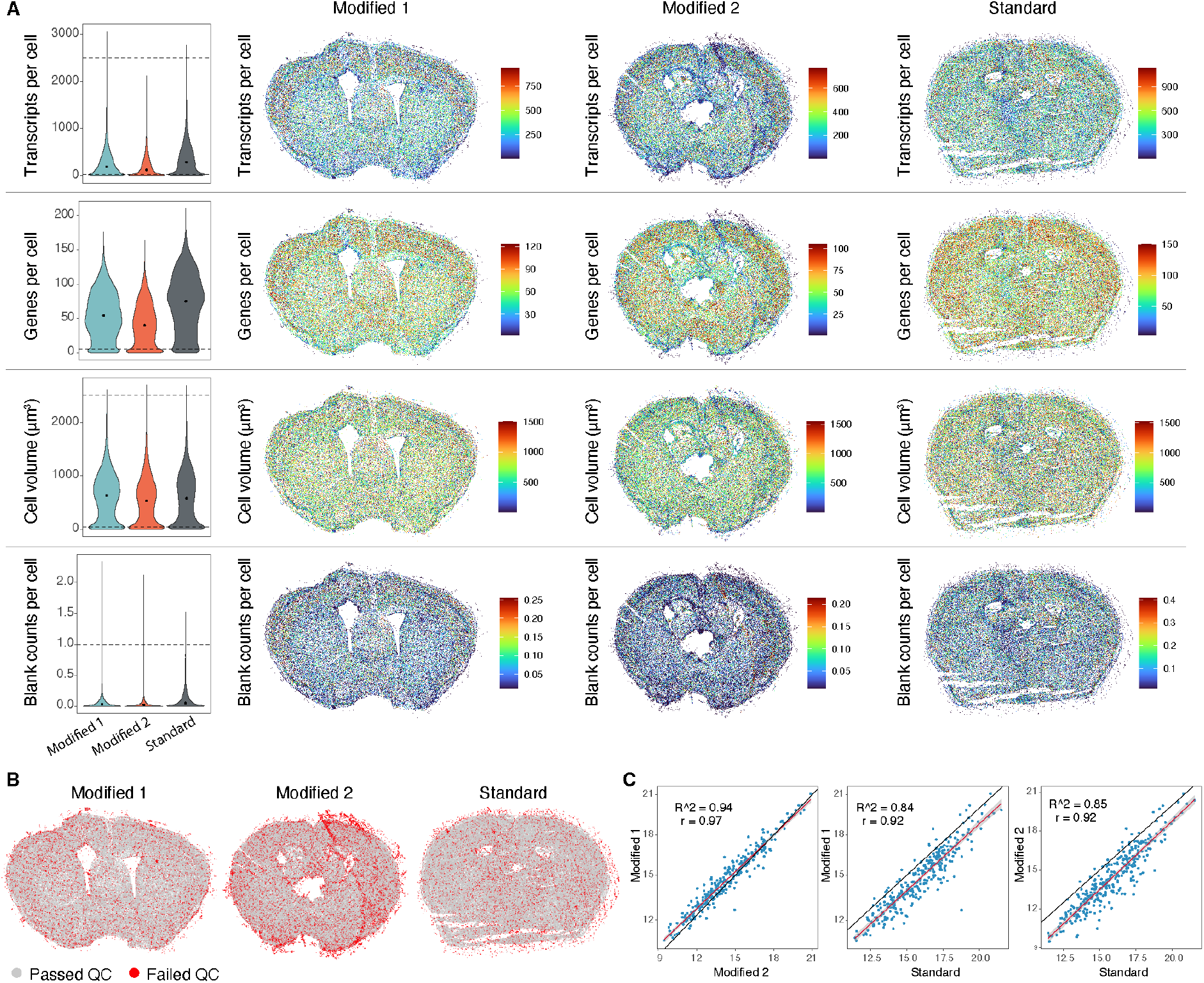
MERFISH quality control (QC) A) QC metrics of MERFISH data in measured brain sections. Dotted lines in violin plots represent cutoff thresholds. Point shows median value. Brain sections on the right show distribution of QC metric values in space. B) Spatial plots showing location of cells passing or failing quality control. C) Correlation of gene expression values between each measured section.

**Figure S2.**
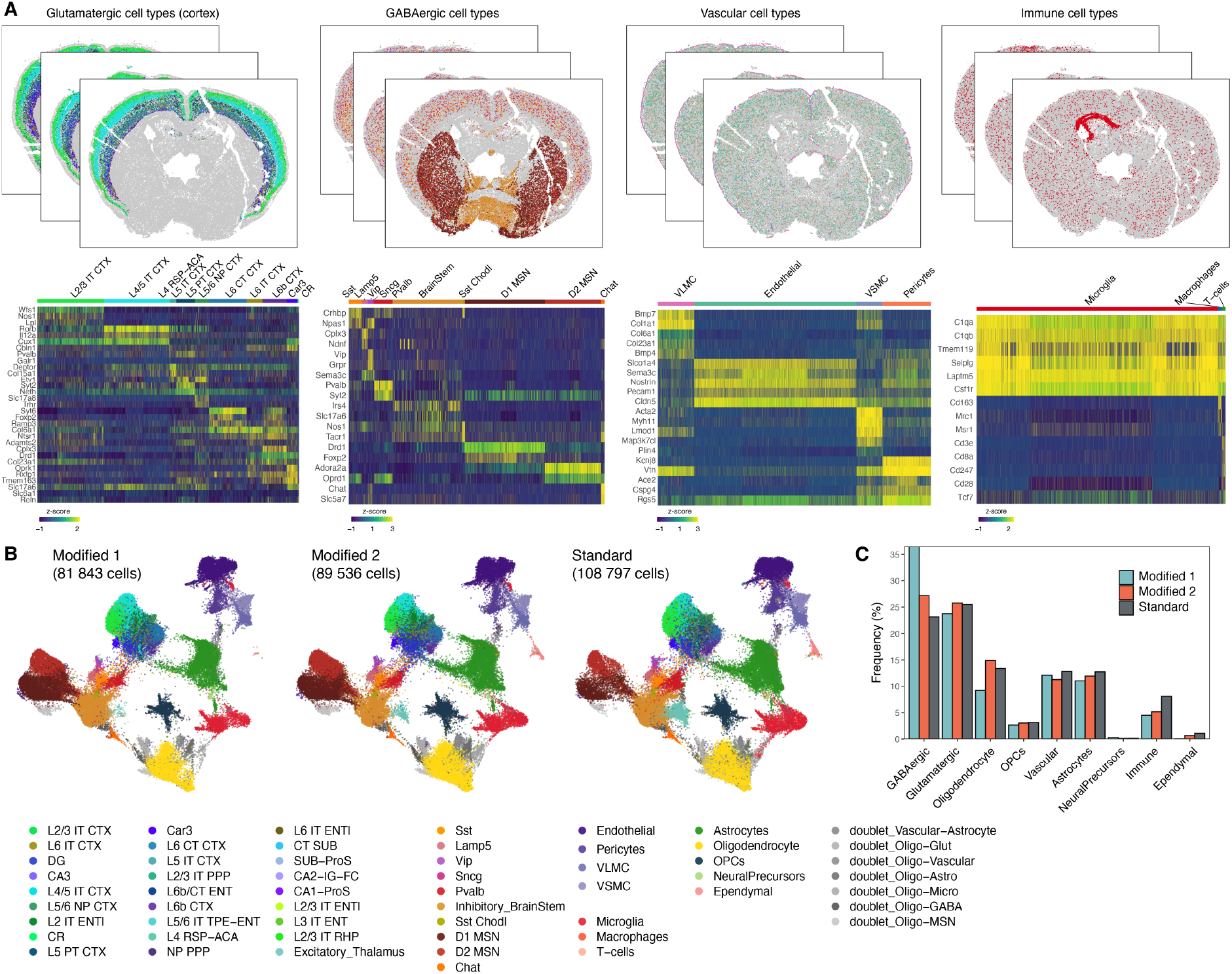
Extended cell type annotation. A) Spatial plots showing location of cell types belonging to major cell classes (top) and expression of their markers (bottom heatmaps). B) UMAP plots split by section and colored by identified cell types. C) Barplot showing frequency of major cell classes per measured section.

**Figure S3.**
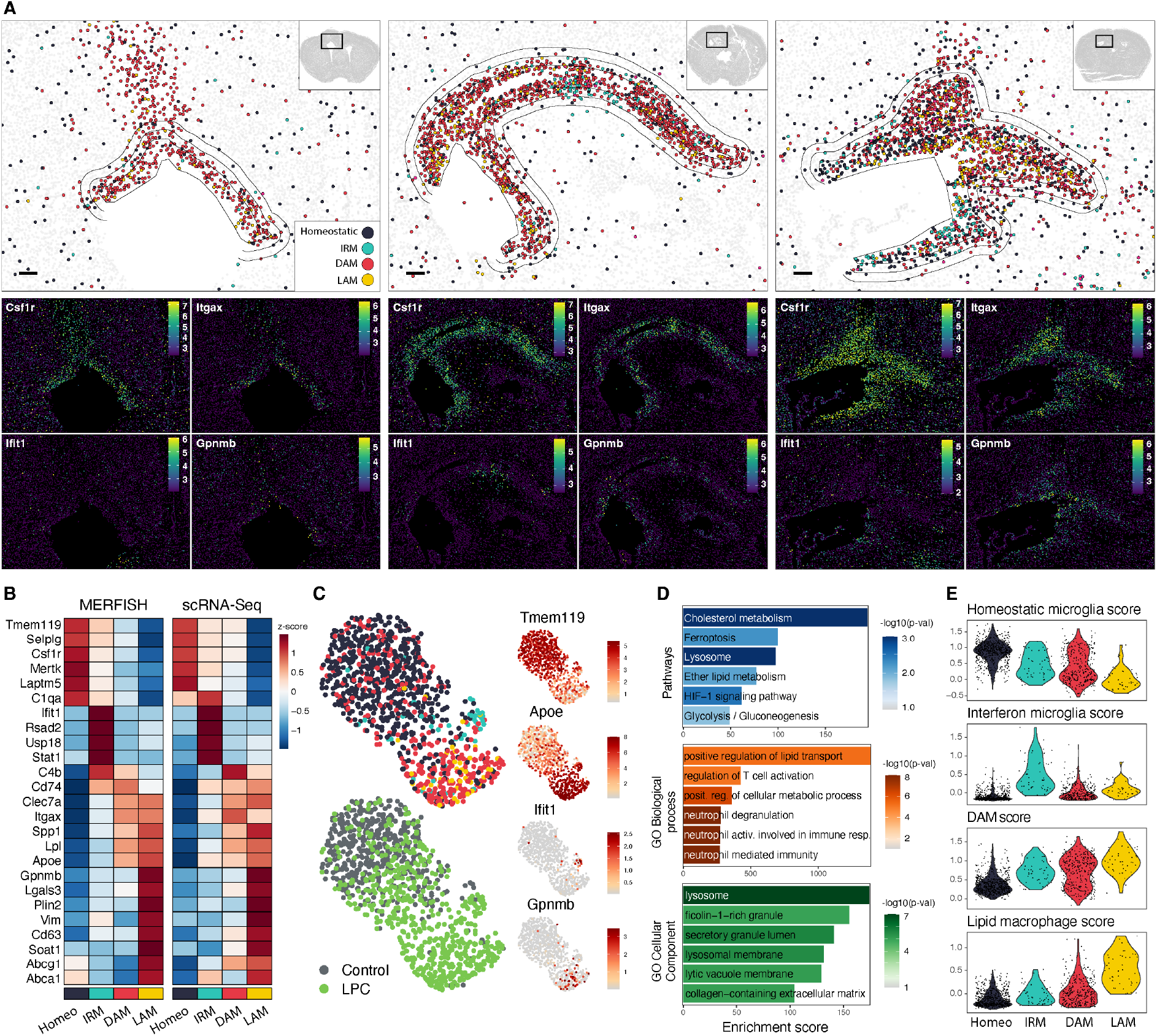
Extended microglia analysis. A) MERFISH spatial plots of microglial clusters in three biological replicate sections (top) and spatial expression of selected marker genes (bottom). Polygons depict segmented lesion areas. Scale bar 100 μm. DAM = disease-associated microglia, IRM = interferon-response microglia, LAM = lipid-associated microglia. B) Heatmap of average expression per microglial cluster measured by MERFISH (left) or by scRNA-Seq (right). C) UMAP embedding of microglial scRNA-Seq data colored by cluster (top left), experimental group (bottom left) and expression of selected markers (right). D) Pathway and Gene Ontology enrichment analysis of identified lipid-associated microglia signature genes. E) Violin plots showing single-cell activity scores of homeostatic microglia, interferon-stimulated microglia, disease-associated microglia and lipid-associated macrophages expression signatures collected from literature.

**Figure S4.**
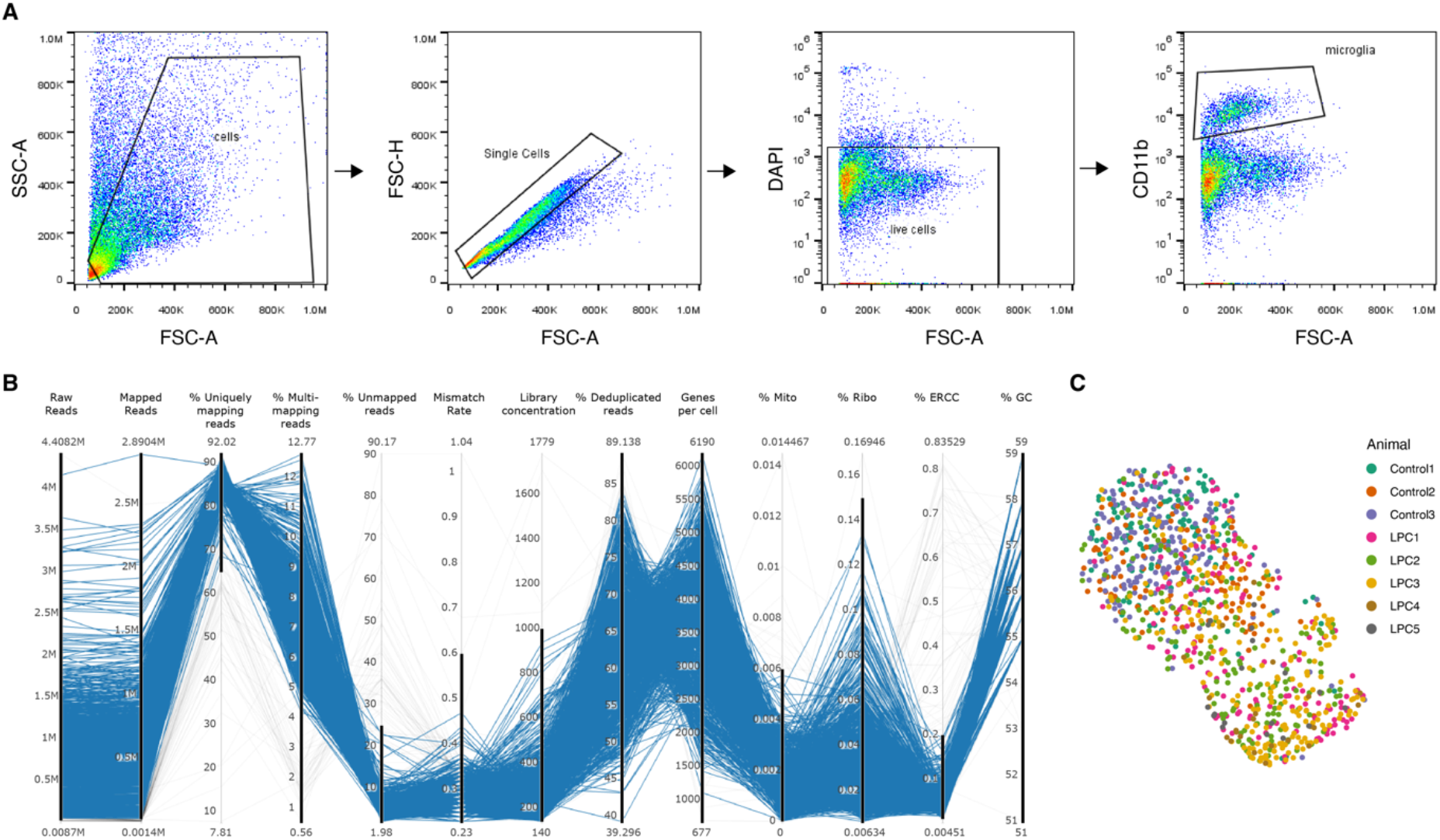
Sorting strategy and quality control of SmartSeq2 data. A) Sorting strategy for SmartSeq2 scRNA-Seq. Flow cytometry gating of CD11b positive cells to enrich microglia. B) Quality control of the SmartSeq2 dataset. Parallel coordinates plot showing cells meeting (blue lines) or failing (grey lines) individual quantitative QC metrics. Bold line segments represent selected threshold boundaries of each QC metric. From left to right, the metrics are: number of raw reads, number of mapped reads, % of uniquely mapping reads, % of multimapping reads, % of unmapped reads, mismatch rate, DNA concentration of final single cell library, % of deduplicated reads, number of detected genes per cell, % of mitochondrial genes, % of ribosomal genes, % of ERCC spike-ins, % of GC content of reads. C) UMAP plot of microglial SmartSeq2 data colored by animal.

## ACKNOWLEDGEMENTS

The work was supported by grants from Chan-Zuckerberg Initiative grant, the German Research Foundation (TRR 128-2, TRR 274/1 – ID 408885537-project Z02, 408885537-408885537. SyNergy Excellence Cluster, EXC2145, Projekt ID390857198), and the Dr. Miriam and Sheldon G. Adelson Medical Research Foundation, and Else Kröner-Fresenius-Stiftung grant. For the single cell and sorting studies, we are grateful for support from the “Flow Cytometric Cell Sorter Sony SH800 Core Unit” run by the Department of Vascular Biology at the Institute for Stroke and Dementia Research (ISD) and SyNergy EXC2145. We thank for technical assistance in EM to Georg Kislinger, Cornelia Niemann, Birgit Kunkel. We also thank for technical assistance with MERFISH to Natalia Petrenko and Jiang He.

## AUTHOR CONTRIBUTIONS

O.G. conceived and supervised the project. P.A., M.S., K.P.A., S.B.-G., H.J., L.C.C., O.G., J.W. L.C., M.S., J.K. planned and performed experiments and analyzed the data, P.A., O.G. wrote the manuscript with input from all authors.

## MATERIALS AND METHODS

### Animals

All mouse experiments in this study were performed with the approval and according to the regulations of the District Government of Upper Bavaria and reported according to guidelines^23, 24^. Male C57BL/6J mice were obtained from Janvier Laboratories. All mice were housed at the animal facility in the German Centre for Neurodegenerative Diseases (DZNE) in Munich in standard, pathogen-free conditions. The temperature in the housing unit was kept between 20 and 22 °C with 40 - 60% humidity and a 12-hour light/12-hour dark cycle.

### LPC injections

Lysophosphatidylcholine (LPC) injections were administered at the age of 16-18 weeks. A solution of 1% LPC (L4129, Sigma) in PBS was mixed with Monastral blue (274011, SigmaAldrich) at a concentration of 0.03% to aid with visualization of the lesion during tissue processing. Mice were anaesthetized with an intraperitoneal injection of MMF solution (0.5 mg medetomidin/kg (body weight), 5.0 mg midazolam/kg (body weight) and 0.05 mg fentanyl/kg (body weight)). Then, head fur was removed, the eyes were treated with bepanthene cream (1578847, Bayer) and a small incision in the skin was performed to expose the skull. The mouse was positioned into a stereotactic injection apparatus and a small hole was drilled at the following injection coordinates (from bregma): X, ± 1.0 mm; Y, −0.1 mm). A glass capillary containing the LPC–monastral blue solution was then lowered to Z: −1.30 mm from bregma, and 1 μL was injected at a rate of 100 nL/minute. Two minutes after the delivery of LPC, the capillary was slowly retracted. The mouse was then injected with 0.05 mg buprenorphin/kg (body weight), and the skin was sutured. Anesthesia was terminated by a subcutaneous injection of AFN solution, containing 2.5 mg/kg (body weight) atipamezol, 1.2 mg/kg (body weight) naloxon and 0.5 mg/kg (body weight) flumazenil.

### Tissue collection and preparation for STcEM

18 days after LPC injection, mice were anaesthetized with an i.p. injection of MMF (fentanyl (0.05 mg/kg)–midazolam (5 mg/kg)–medetomidine (1 mg/kg)) and transcardially perfused with 2 UI/mL Heparin (Heparin-Natrium-25000-ratiopharm^®^, PZN: 03029843) in HBSS (no calcium, no magnesium, Gibco™, 14175129) for 3 min and 4 % paraformaldehyde (PFA, EM Grade, Electron Microscopy Sciences, Cat. No. 15710, diluted in 10X PBS, Invitrogen™, AM9624 and UltraPure™ Distilled Water, Invitrogen™, 10977-035) for 5 min, before carefully removing the brain from the skull. Afterwards, the brains were fixed by submergingin 4% PFA for 6 hours, followed by 14 hours in 15% sucrose (Sigma-Aldrich, S0389 in UltraPure™ Distilled Water, Invitrogen™, 10977-035) and 5 hours in 30% sucrose. Next, the PFA-fixed brains were simultaneously embedded in Tissue-Tek^®^ O.C.T.™ Compound (Sakura, 4583) and frozen in a plastic mold on dry ice. For the fresh frozen brain samples, the mouse was only perfused with Heparin in HBSS and directly embedded and frozen in Tissue-Tek^®^ O.C.T.™ Compound on dry ice (“standard protocol”). Brains were stored at −80°C until further processing.

### Tissue sectioning

Coronal, 10 μm thick brain sections were prepared and collected at a cryotome (CryoStar NX70, Thermo Scientific). Sections determined for MERFISH analysis were placed on round glass slides provided by Vizgen Corp. (Cambridge, MA 02138; MERSCOPE slide part number 20400001). Sections for electron microscopy were collected on Superfrost Plus^®^ Gold slides (Thermo Scientific, Menzel Gläser, K5800AMNZ72). Brain sections for MERFISH were subsequently washed two times with PBS and one time with 70 % ethanol (VWR Chemicals, 20.821.310, diluted in UltraPure™ Distilled Water, Invitrogen™) for 5 minutes each. Then samples were individually sealed in bags filled with 70 % ethanol and shipped to Vizgen Corp. (Cambridge, MA 02138) for MERFISH analysis. Sections for electron microscopy were stored at −80°C until further processing.

### MERFISH procedure

#### Gene panel

Gene panel for this study consisted of 287 protein-coding genes and 98 blank probes. Genes included selection of known brain and immune cell type markers including of glial cells, T-cells, macrophages and subtypes of Glutamatergic and GABAergic neurons. In addition we included a panel of microglial, astrocytic and oligodendrocytic reactive markers from literature, and genes from cholesterol metabolic pathways. Full gene panel is listed in supplementary file.

#### Hybridization

Samples on beaded slides were placed tissue-side up in 60×15 mm petri dishes and kept at the back of the cryostat at −20° C for at least 5 minutes for the tissue to adhere. For the fresh frozen samples, 5 ml of fixation buffer (4% paraformaldehyde in 1x Phosphate buffered saline) were added in a fume hood and incubated for 15 min at room temperature. The fresh frozen samples were then washed with 5ml Phosphate buffered saline 3 times, 5 minutes each.

Both the fresh frozen and the fixed frozen samples were then permeabilized in 5ml 70% ethanol at 4° C overnight, in parafilm-sealed dishes, and stored long-term in the same conditions. For hybridizing with the library (the gene panel), the samples were washed with 5ml Vizgen Sample Prep Wash Buffer (Vizgen part number 20300001) and then incubated in 5ml Formamide Wash Buffer (Vizgen pn 20300002) at 37° C for 30 min in an incubator. The Formamide Wash Buffer was aspirated from the tissue, and 50 μl of the gene panel mix was added on top of each tissue. A piece of parafilm ~1.5cm x 1.5cm was placed on top to spread the library mix and protect it from evaporation. The dishes were sealed with parafilm and placed in a humidified incubator at 37° C for 36-48 hours. The parafilm was removed from the top of each tissue, and the samples were incubated in 5ml Formamide Wash Buffer at 47° C for 30 minutes, twice. The samples were then washed with 5ml Sample Prep Wash Buffer for 2 minutes.

#### Gel embedding

To gel embed the samples, fresh 10% w/v ammonium persulfate solution was prepared. For each sample, 5ml of Gel Embedding Premix (Vizgen pn 20300004) was combined with 25 μl of the 10% ammonium persulfate solution and 2.5 μl of TEMED (N,N,N’,N’-tetramethylethylenediamine). In parallel, one 20mm Gel Coverslip (Vizgen pn 20400003) for each sample was cleaned with RNAseZap, followed by 70% ethanol and dried with Kimwipes. The Gel Coverslips were then covered with 100 μl of Gel Slick Solution (VWR, catalog number 12001-812) for a minute and wiped dry with Kimwipes. The Sample Prep Wash Buffer was aspirated from the samples. For each sample, 100 μl of the Gel Embedding Mix was retained in a small tube, while the remainder of the Gel Embedding Mix was added to the samples and incubated for 1 minute. The Gel Embedding Mix was then poured out from the samples into a waste tube but kept aside on the bench (to monitor gel formation). The slides were then aspirated dry, leaving just enough liquid to keep the tissue from drying out. 85 μl of the separately retained Gel Embedding Mix was added on top of the tissue, and the Gel Slick treated coverslip was placed over it with tweezers, with the Gel Slick-treated side facing down toward the tissue and avoiding air bubbles. Extra Gel Embedding Solution was aspirated from the sides of the coverslips. The dishes were incubated at room temperature for 1.5 hours to allow the gels to form. Thereupon, the coverslips were removed using a Hobby Blade and tweezers.

#### Tissue clearing

To clear the samples of lipids and proteins that interfere with imaging, 5 ml of Clearing Premix (Vizgen pn 20300003) were mixed with 50 μl of Proteinase K for each sample. After the coverslips were removed from the gel embedded samples, the clearing solution was added to each sample, and the dishes were sealed with parafilm. The fresh frozen samples were placed at 37° C in a humidified incubator overnight, while the fixed frozen samples were placed at 47° C in a humidified incubator overnight (or for a maximum of 24 hours), and then moved to 37° C. The samples were stored in the Clearing solution in the 37° C incubator prior to imaging for up to a week.

#### Sample imaging

The Clearing solution was aspirated from the sample, and the sample was washed three times with Sample Prep Wash Buffer briefly, then again for 10 minutes on a rocker, and then three more times briefly. The sample was incubated with 3 ml of the appropriate first hybridization buffer, including Dapi and polyT reagent (Vizgen pn 20300021), for 15 minutes at room temperature on a rocker, covered from light. The sample was then washed with 5ml of the Formamide Wash Buffer (Vizgen pn 20300002) for 10 minutes at room temperature on a rocker, covered from light, and then transferred to 5ml of the Sample Prep Wash Buffer (Vizgen pn 20300001). In the meantime, the Imaging buffer was prepared by combining the Imaging buffer, the Imaging Buffer Activator (Vizgen pn 20300015), and RNase inhibitor at a ratio of 500: 2.5: 1. The hybridization buffers appropriate to the gene panel, as well as the imaging buffers, were loaded onto the Vizgen microscope system. The sample was placed in the flow chamber and connected to the fluidics system of the Vizgen microscope, taking care to disperse air bubbles. A low-resolution mosaic was acquired using a 10X objective, and the regions of interest were selected for high-resolution imaging with a 60x lens.

For the high resolution imaging, the focus was locked to the fiducial fluorescent beads on the coverslip. Seven 1.5 μm-thick z planes were taken for each field of view when imaging the tissue, including for the DAPI channel. Cell segmentation was performed using the Watershed algorithm, using DAPI nuclear seeds and PolyT total RNA staining basins. Images were decoded to RNA spots with xyz and gene id using Vizgen’s Merlin software.

### MERFISH Analysis

#### Data quality control and filtering

Single-cell gene expression matrices were obtained by counting mRNA molecules within segmented cell boundaries and further analyzed in R using Seurat package and custom-made scripts. We excluded cells containing less than 30 or more than 2500 individual transcripts, less than 5 unique genes, cells with volume less than 40 μm^3^ or more than 2500 μm^3^ or cells with average count of blank probe spots more than 1.

#### Annotation of cell types and clustering analysis

Data were normalized by dividing gene counts for each cell by total count for that cell, multiplied by 10 000 and log-transformed. Data were then scaled and principal components were calculated on all 287 measured genes. Between X and Y principal components were used to calculate UMAP embedding and perform clustering analysis using Louvain algorithm. After examining each section individually, we integrated data from three replicates using Seurat’s rpca workflow and repeated UMAP and clustering analysis. Cluster markers were identified using Wilcoxon rank-sum test. To resolve neuronal subclasses, we downloaded Allen Brain Atlas reference single cell RNA-Seq data, including metadata with cell type annotations^25^. We then substracted the reference data to neuronal classess and mapped cell type annotations to our integrated MERFISH data using Seurat’s label transfer workflow. These annotations as well as annotation of remaining cell types (striatal neurons, glial, immune cells) were refined in rounds of subclustering analysis by each time subsetting data to major cell class and repeating normalization, scaling, pca and clustering workflows. During analysis we noted that fraction of segmented single cell profiles exhibits contamination with transcripts from other cell types originating mostly from imperfect cell boundary segmentation. These resemble “doublets”, however unlike doublets in droplet-based single-cell RNA-Seq workflows, in MERFISH data they bear biological significance as they originate from physically proximal cells. Therefore, we opted to annotate clusters that clearly represented mixture of different cell types as doublets, but keep them in the data, keeping this in mind during analyses. During subclustering analysis, we further removed effect of contaminating RNA by regressing expression signatures of other cell types from the data. For this, we first obtained signatures of each cell type by searching for differentially expressed genes between analyzed cluster and other coarse clusters (other present cell types) using strict threshold. We then calculated aggregate score for each signature in each cell using Seurat’s AddModuleScore function and regressed these scores from expression matrix of analyzed cluster before running PCA, UMAP and subclustering analyses. For subclustering of microglia, we further excluded known non-microglial genes (known markers of other cells) from the matrix. This strategy dramatically improved resolution of clusters and allowed us to discover expression patterns previously masked by contaminating RNA while keeping sufficient cell numbers.

#### Spatial analysis of the lesion

To quantify cell types within the lesion areas, we in silico dissected lesion areas into lesion core, lesion inner edge and lesion outer edge. First, we segmented area representing lesion core based on: i) spatial pattern of microglia (accumulated in lesion core, decreasing density towards lesion edge), ii) spatial pattern of oligodendrocytes (absent in lesion, marking lesion edge) and iii) expression profile of Mbp (marking lesion edge). We then expanded the polygon around lesion core twice by 50 um, segmenting inner lesion edge and outer lesion edge. For comparison, we also segmented area of uninjured white matter and uninjured cortical grey matter from contralateral hemisphere. We then quantified proportions of cell types in these areas.

#### Cell neighborhood analysis

To analyze cellular neighborhoods of microglial clusters and T-cells in the lesion, we first subsetted data to union of cells from lesion core, lesion inner edge and lesion outer edge. We then identified each cell’s 5 nearest neighbors in physical space (based on Euclidean distance) and counted the proportion of each cell type label among the neighbors. We then repeated this process 10 000 times, each time randomly permuting cell type labels to obtain empirical p-value and empirical fold-enrichment (defined as observed fraction of cell type label among neighbors divided by average fraction obtained from all permuted iterations). Neighborhood analysis of the EM data was performed in the same way using EM-derived annotations and cellular positions.

#### Alignment of MERFISH and EM data

DAPI staining images of MERFISH sections were registered onto EM overview scans of the sections of the same sample based on user-defined anatomical landmarks with BigWarp ImageJ plugin using thin plate spline transformation.

### Tissue collection and preparation for scRNA-Seq

The mice were deeply anesthetized and perfused with cold HBSS between 9am-11am (to decrease circadian fluctuations). Each brain was removed and under a dissection microscope individually micro-dissected; gray matter was isolated from the frontal cortex and white matter form optic tract, medial lemniscus and corpus callosum (attached gray matter and choroid plexus were carefully removed). We used a microglia isolation protocol we previously described^26^, that prevents ex-vivo transcription and automatizes the mechanical isolation parts using GentleMacs with the Neural Tissue Dissociation Kit (Papain) (Miltenyi Biotec). We added actinomycin D (Act-D, Sigma, No. A1410) to a final concentration of 45 μM into the dissociation solution and enzyme mix to prevent ex-vivo transcription. The dissociated cell suspension was passed through a 70 μm cell strainer (Corning, 352350) before labeling. Subsequently, cells were blocked with mouse FcR-blocking reagent (CD16/CD32 Monoclonal Antibody, eBioscience cat:14-0161-82,1100) and then stained with the antibody against CD11b (PE/Cy7,M1/70, eBioscience, Cat:48-0451-82,1:200) and washed with PBS (Sigma, D8537). Then the cells were then stained with DAPI (4’,6-diamidino-2-phenylindole, 1:10000 dilution; Sigma) to label dead cells. Viable (DAPI negative) single immune cells (CD11b positive cells) were sorted by flow cytometry (SH800; Sony). Single-cells were sorted into 96 well plates filled with 4 μL lysis buffer containing 0.05% Triton X-100 (Sigma), ERCC (External RNA Controls Consortium) RNA spike-in Mix (Ambion, Life Technologies) (1:24000000 dilution), 2.5 μM oligo-dT, 2.5 mM dNTP and 2 U/μL of recombinant RNase inhibitor (Clontech) then spun down and frozen at −80° C.

### scRNA-Seq library preparation and sequencing

The 96-well plates containing the sorted single cells were first thawed and then incubated for 3 min at 72°C and thereafter immediately placed on ice. To perform reverse transcription (RT) we added into each well a master mix of 0.59 μL H2O, 0.5 μL SMARTScribe™ Reverse Transcriptase (Clontech), 2 μL 5x First Strand buffer, 0.25 μL Recombinant RNase Inhibitor (Clontech), 2 μL Betaine (5 M Sigma), 0.5 μL DTT (100 mM) 0.06 μL MgCl2 (1 M Sigma), 0.1 μL

Templateswitching oligos (TSO) (100 μM AAGCAGTGGTATCAACGCAGAGTACrGrG+G). Next RT reactions were incubated at 42°C for 90 min followed by 70°C for 5 min and 10 cycles of 50°C 2 min, 42°C 2 min; ending with 70°C for 5 min for enzyme inactivation. Preamplification of cDNA was performed by adding 12.5 μL KAPA HiFi Hotstart 2x (KAPA Biosystems), 2.138 μL H2O, 0.25 μL ISPCR primers (10 μM, 5’ AAGCAGTGGTATCAACGCAGAGT-3), 0.1125 μL Lambda Exonuclease under the following conditions: 37°C for 30 min, 95°C for 3 min, 23 cycles of (98°C for 20 sec, 67°C for 15 sec, 72°C for 4 min), and a final extension at 72°C for 5 min. Libraries were then cleaned using AMPure beads (Beckman-Coulter) cleanup at a 0.7:1 ratio of beads to PCR product. Libraries were assessed by Bio-analyzer (Agilent 2100), using the High Sensitivity DNA analysis kit, and quantified using Qubit’s DNA HS assay kits and a Qubit 4.0 Fluorometer (Invitrogen, LifeTechnologies). Further selection of samples was performed via qPCR assay against ubiquitin transcripts Ubb77 (primer 1 5’-GGAGAGTCCATCGTGGTTATTT-3’ primer 2 5’-ACCTCTAGGGTGATGGTCTT-3’, probe 5’-/5Cy5/TGCAGATCTTCGTGAAGACCTGAC/3IAbRQSp/-3’) measured on a LightCycler 480 Instrument II (Roche). Samples were normalized to 160 pg/μL. Sequencing libraries were constructed by using in-house produced Tn5 transposase. Libraries were barcoded, pooled and purified in 3 rounds of AMPure bead (Beckman-Coulter) cleanup at a 0.8:1 ratio of beads to library. Libraries were then sequenced with 100 bp paired-end sequencing on DNBSeq platform (BGI group) to a median depth of 8.6×10^5^ reads/sample.

### scRNA-Seq analysis

Demultiplexed Fastq files were quality-controlled with FastQC and reads were then aligned using rnaSTAR to the GRCm38 (mm10) genome with addition of ERCC spike-in sequences. To obtain single cell gene expression matrices reads were counted with rnaSTAR using parameter “quantMode GeneCounts” and unstranded argument. Further analysis was performed in R using Seurat package and custom-made scripts. Samples were filtered for quality with several QC thresholds (Figure S4b). Data from LPC-injected and control mice were integrated together using Seurat’s CCA integration workflow: Expression was normalized by dividing gene counts for each cell by total count for that cell, multiplied by 10 000 and log-transformed, 3000 variable features were identified with SelectIntegrationFeatures() function, transfer anchors were found using FindIntegrationAnchors() function and data were integrated with IntegrateData() function. Data were scaled before calculating PCA and UMAP embedding using 30 PCs. Cell type clusters were identified using Leiden algorithm and annotated based on canonical cell type markers identified by Wilcoxon rank-sum test. After first round of coarse clustering, microglia were isolated and analyzed separately by repeating aforementioned workflow. Microglial cluster labels were then mapped from MERFISH data onto SmartSeq2 data by finding anchors using Seurat’s FindTransferAnchors() function and mapping labels with TransferData() functions. Cluster markers were identified using Wilcoxon rank-sum test.

#### Functional enrichment analysis

Gene expression signatures for each microglial population were identified by i) finding sets of differentially upregulated genes between said population and every other population (wilcoxon rank-sum test) ii) intersecting these sets. Identified signatures are available in supplementary file. Enriched KEGG pathways and Gene Ontology terms in these signatures were then identified using Enrichr package. Full results are available in supplementary file. For analysis of published microglial signatures, marker genes were collected from relevant publications. Specifically, for lipid-associated macrophage signature we intersected markers of lipid-associated macrophages in adipose tissue^15^, atherosclerotic aorta^16^ and liver^17^. For DAM signature, we intersected markers of disease-associated microglia (DAM)^11^, activated-response microglia (ARM)^12^, and neurodegeneration-related microglia signature^27^ and excluded genes in lipid-associated macrophage signature to assure specificity. For homeostatic signature we intersected markers of homeostatic microglia from^11, 27^. For interferon signature we intersected signature of interferon microglia^27^ and core module of interferon-stimulated genes^28^. Activity of each signature was scored in each cell using AddModuleScore() function. Signatures from literature are available in supplementary file.

### Electron microscopy

#### Serial section electron microscopy using automated tape-collecting ultramicrotomy (ATUM)

Mouse cryotome sections adjacent to the ones analyzed by spatial transcriptomics were sectioned at 10 μm thickness and collected onto glass slides (SuperFrost Plus Gold, Thermo). Tissue sections stayed adherent to the glass slides during the processing for EM and were kept in slide containers (Simport™ Scientific LockMailer™ Tamper Evident Slide Mailer, Fisher Scientific). These holders were positioned in a wrack on an orbital shaker during the incubation steps and reagents were exchanged by pouring or pipetting. We postfixed the sections in 2.5% glutaraldehyde (Science Services) in 0.1 M sodium cacodylate (Science Services) buffer (pH 7.4) for 30 min and applied a standard rOTO protocol starting with 1h incubation in 1% osmium tetroxide (Science Services), 1% potassium ferricyanide (Sigma) in 0.1 M sodium cacodylate buffer (pH 7.4). After washing and reaction with 1% thiocarbohydrazide (Sigma) for 20 min at 40°C we applied a second osmium step (1% osmiumtetroxide in water, 1h). The tissue was further contrasted in 1% aqueous uranyl acetate at 4°C over night. Samples were dehydrated in an ascending ethanol series and infiltrated with LX112 in acetone (LADD). Large gelatin capsules covering the entire brain coronal section (size 000, 9.55 mm diameter, Science Services) were positioned onto the glass slides and cured for 2d at 60°C. In order to remove the encapsuled sample from the glass slide we notched the resin around the tissue section with a sharp blade. The glass slide was then submerged into liquid nitrogen for several seconds and heated up in a 60°C water bath. These freeze-thaw cycles were repeated until the tissue block could be removed from the glass slide.

The block was trimmed at the empty resin end using a rotary tool (Dremel) in order to fit it into a standard sample holder. We trimmed the tissue end to generate an approximately 3×5mm block face bearing the entire cortex and the ventriclesusing a trimming machine (TRIM2, Leica). Serial sections at 150-200 nm thickness were taken on an ATUMtome (Powertome, RMC) using a histo knife (Diatome) and collected on freshly plasma-treated (custom-built, based on Pelco easiGlow, adopted from M. Terasaki, U. Connecticut, CT), carbon nanotube (CNT) tape (Science Services). CNT tape stripes were assembled onto adhesive carbon tape (Science Services) attached to a 4-inch silicon wafer (Siegert Wafer) and grounded by adhesive carbon tape strips (Science Services). EM micrographs were acquired on a Crossbeam Gemini 340 SEM (Zeiss) with a four-quadrant backscatter detector at 8 kV. In ATLAS5 Array Tomography (Fibics), we acquired the whole section at 200 nm and ipsi- and contralateral regions of interest at 20 nm lateral resolution. We selected single regions for acquisition at high resolution (4 nm pixel size). Serial section data were stitched, aligned and analyzed in Fiji TrakEM2^29^.

For the whole area annotation (Fig. 2) we trimmed the block face to a 2×3 mm size covering the ipsilateral lesion site. Serial sections (127 at 200 nm thickness) were taken and collected onto tape. We imaged the whole section overview at 200×200×200 nm resolution and selected the region of interest within a volume of 13.2 μm (66×200 nm) thickness. Every second section (z resolution 400 nm) was imaged at 20×20 nm lateral resolution. This resulted in three image stacks, one covering the full area of interest at 1.2×1.2 mm and two covering 0.5×0.5 mm. The large image stacks were exported as tiles, stitched using TrakEM2 and three VAST files generated from them. Annotation of cell types according to the ultrastructural morphology was performed in VAST. The respective cell of interest was investigated along the entire stack thickness, screened for ultrastructural features (lipid droplet, lysosomal content, ER branching), categorized and flagged at the stack surface. The three VAST object files were exported and reassembled in Blender.

